# BrnQ-type Branched-chain Amino Acid Transporters Influence *Bacillus anthracis* Growth and Virulence

**DOI:** 10.1101/2021.12.09.472043

**Authors:** Soumita Dutta, Ileana D. Corsi, Naomi Bier, Theresa M. Koehler

**Author notes:** Correspondence Theresa M. Koehler.

## Abstract

*Bacillus anthracis,* the anthrax agent, exhibits robust proliferation in diverse niches of mammalian hosts. Metabolic attributes of *B. anthracis* that permit rapid growth in multiple mammalian tissues have not been established. We posit that branched-chain amino acid (BCAA: Isoleucine, leucine and valine) metabolism is key to *B. anthracis* pathogenesis. Increasing evidence indicates relationships between *B. anthracis* virulence and expression of BCAA-related genes. Expression of some BCAA-related genes is altered during culture in bovine blood *in vitro* and the bacterium exhibits valine auxotrophy in a blood serum mimic medium. Transcriptome analyses have revealed that the virulence regulator AtxA, that positively affects expression of the anthrax toxin and capsule genes, negatively regulates genes predicted to be associated with BCAA biosynthesis and transport. Here, we show that *B. anthracis* growth in defined media is severely restricted in the absence of exogenous BCAAs, indicating that BCAA transport is required for optimal growth *in vitro*. We demonstrate functional redundancy among multiple BrnQ-type BCAA transporters. Three transporters are associated with isoleucine and valine transport, and deletion of one, BrnQ3, attenuates virulence in a murine model for anthrax. Interestingly, an *ilvD*-null mutant lacking dihydroxy-acid dehydratase, an enzyme essential for BCAAs synthesis, exhibits unperturbed growth when cultured in media containing BCAAs, but is highly attenuated in the murine model. Finally, our data show that BCAAs enhance AtxA activity in a dose-dependent manner, suggesting a model in which BCAAs serve as a signal for virulence gene expression.

**IMPORTANCE:** Infection with *B. anthracis* can result in systemic disease with large numbers of the bacterium in multiple tissues. We found that BCAA synthesis is insufficient for robust growth of *B. anthracis*; access to branched-chain amino acids (BCAAs) is necessary for proliferation of the pathogen during culture and during infection in a murine model for anthrax. *B. anthracis* produces an unusually large repertoire of BCAA-related transporters. We identified three isoleucine/valine transporters with partial functional redundancy during culture. Deletion of one of these transporters, BrnQ3, resulted in attenuated virulence. Interestingly, a BCAA biosynthesis mutant grew well in medium containing BCAAs, but like BrnQ3, was attenuated for virulence. These results suggest that BCAAs are limiting in multiple niches during infection and furthers understanding of nutritional requirements of this important pathogen.

## INTRODUCTION

*Bacillus anthracis*, the bacterium that causes anthrax, is well known for its robust proliferation in diverse niches of mammalian hosts. Infection can result in up to 10^8^ colony-forming units (CFU) per gram of tissue, including various organs, blood, and cerebral spinal fluid, at the time of host death (1–5). While the roles of several secreted virulence factors, including the anthrax toxin proteins, poly-D-glutamic acid capsule, siderophores, and proteases, have been discerned for anthrax (2, 6–9), the metabolic attributes of *B. anthracis* that permit rapid proliferation to high numbers in multiple mammalian tissues have not been established. *B. anthracis* is a facultative anaerobe and grows in most rich undefined media with a doubling time of approximately 30 minutes. Defined minimal media that support *B. anthracis* growth *in vitro* contain glucose, salts, and nine or more amino acids, but reports differ regarding essential nutrients (10, 11). The *B. anthracis* genome reveals a repertoire of metabolic and transport genes that are homologous to genes of the well-studied and nonpathogenic *Bacillus* species, *B. subtilis* (12, 13). Yet remarkably, genomic and transcriptomic analyses indicate that *B. anthracis* has a large capacity for amino acid and peptide utilization, compared to *B. subtilis* (4, 12, 14, 15). Here we explore branched-chain amino acid (BCAA) metabolism by *B. anthracis* as a potential key aspect of anthrax pathogenesis.

The BCAAs, isoleucine, leucine, and valine, are required for protein synthesis and serve as precursors of branched-chain fatty acids, the major fatty acids of the Gram-positive bacterial cell membrane (16). BCAAs can also act as signals related to nutritional status. For many low G+C Gram-positive bacteria, BCAAs are effectors for the global transcriptional regulator CodY, which modulates gene expression to support environmental adaptation (17, 18). In some pathogens, BCAA biosynthesis, BCAA transport, or both processes, can affect virulence (16, 19–22). BCAAs, like all amino acids, can be synthesized from intermediates of central metabolic pathways if the bacterium possesses the appropriate biosynthetic enzymes, or can be transported into the bacterial cell from the environment if the cell envelope includes proper transport machinery.

The BCAA biosynthesis pathway is highly conserved among bacteria (23). The pathway is composed of eight enzymes, some of which are involved in biosynthesis of multiple BCAAs and some of which are BCAA-specific. Interestingly, despite possessing all enzymes required for BCAA biosynthesis, culture of some bacteria, including *Listeria monocytogenes*, *Streptococcus pneumoniae*, and *Streptococcus suis*, requires the presence of BCAAs (24–26). Others, such as *Staphylococcus aureus*, show significantly delayed growth in media deficient for one or more BCAAs (27).

Acquisition of BCAAs from the environment can be accomplished by some bacteria that use specialized transporters to bring BCAAs into the cell cytosol. Putative BCAA transporter genes are commonly found in bacterial genomes; however, transporters have only been characterized in a few species (16, 28–30). BCAA transport has been described most extensively in *B. subtilis, Lactococcus lactis*, and in the pathogen *S. aureus*. In *B. subtilis,* BrnQ, BcaP, and BraB are major transporters for isoleucine and valine, and are likely also able to transport leucine (29). In *S. aureus*, BrnQ1, BrnQ2, and BcaP have been characterized as BCAA transporters (16, 21). Two BCAA transporters with redundant function have been reported for *L. lactis* (31, 32), while *Corynebacterium glutamicum* and *Lactobacillus delbrueckii* appear to each carry only one BCAA transporter (28, 30).

BCAA synthesis and transport have not been well-studied in *B. anthracis*, however some reports suggest that BCAA metabolism is related to *B. anthracis* pathogenesis. *B. anthracis* exhibits valine auxotrophy when cultured in a blood serum mimic medium (BSM) (33). When *B. anthracis* is grown in bovine blood *in vitro*, genes for putative BrnQ-related transporters are highly induced, whereas BCAA-biosynthesis related genes and other BCAA-associated ABC transporter genes are repressed (34). Perhaps most intriguingly, transcript levels of *brnQ*-related transport genes and predicted BCAA biosynthesis genes respond to host-related cues during culture and are regulated by AtxA, a critical regulator of virulence in *B. anthracis.* AtxA positively affects expression of the anthrax toxin and capsule genes (35–37) and is essential for virulence in some animal models for anthrax (36, 38). Recently, Leppla and coworkers demonstrated specific binding of AtxA to the *pagA* promoter region (41). While the precise mechanism for AtxA function is not clear, we have demonstrated that AtxA activity and multimerization is enhanced by host-related signals including CO_2_/bicarbonate and glucose (5, 39, 40). Interestingly, our RNA-seq data reveal that AtxA strongly represses the expression of BCAA transport and biosynthesis genes (15), and the regulation is possibly mediated by the AtxA-controlled small regulatory RNA XrrA (4).

Here we describe the genomic arrangement of BCAA-related genes in *B. anthracis* (https://www.ncbi.nlm.nih.gov/nucleotide/AE017334). BCAA biosynthesis genes are clustered into two operons, while BrnQ-related BCAA transporter genes are present in multiple loci. We assess the function and specificity of putative BrnQ BCAA transporters and examine relationships between BCAAs and AtxA. Finally, we test the virulence of mutants disrupted for BCAA transport and biosynthesis in a murine model for systemic infection. Our data indicate that both BCAA transport and synthesis are required for *B. anthracis* virulence, establishing a connection between central metabolism and pathogenesis in *B. anthracis*.

## RESULTS

### Growth of *B. anthracis* in toxin-inducing conditions with and without BCAAs

Previous studies have shown that *B. anthracis* growth in various media is largely affected by isoleucine, leucine and/or valine (33, 42, 43). We wanted to test the requirement for BCAAs when *B. anthracis* is cultured in a defined medium in conditions known to promote synthesis of the anthrax toxin proteins. R medium (10) was formulated based on the content of a casamino acids (CA) medium that was used previously to promote toxin synthesis (44, 45). R medium contains defined concentrations of glucose, salts, amino acids, vitamins and nucleotides. Toxin synthesis is maximized when *B. anthracis* is cultured in CA (undefined) or R (defined) containing 0.8% bicarbonate and incubated at 37°C, with shaking in a 5% CO_2_ atmosphere (10, 11, 38, 45).

The BCAA content of R medium is 1.75 mM isoleucine, 1.50 mM leucine, and 1.35 mM valine (11). As shown in Fig. 1A, *B. anthracis* grown in R medium in toxin-inducing conditions exhibited a doubling time of approximately 1 h and a final OD_600_ of about 1. The absence of all three BCAAs resulted minimal growth of the bacterium during 20 h of incubation in these conditions. Further, culture in R medium in the absence of individual BCAAs revealed that leucine and valine are critical for *B. anthracis* growth. Poor growth in the absence of these amino acids suggests that *B. anthracis* relies on transport, rather than biosynthesis of leucine and valine for optimal growth. By contrast, the absence of isoleucine had a relatively small but reproducible adverse effect on growth, indicating that isoleucine biosynthesis is sufficient for growth in these conditions, but transport of the amino acid offers a small growth advantage.

**Figure 1:**
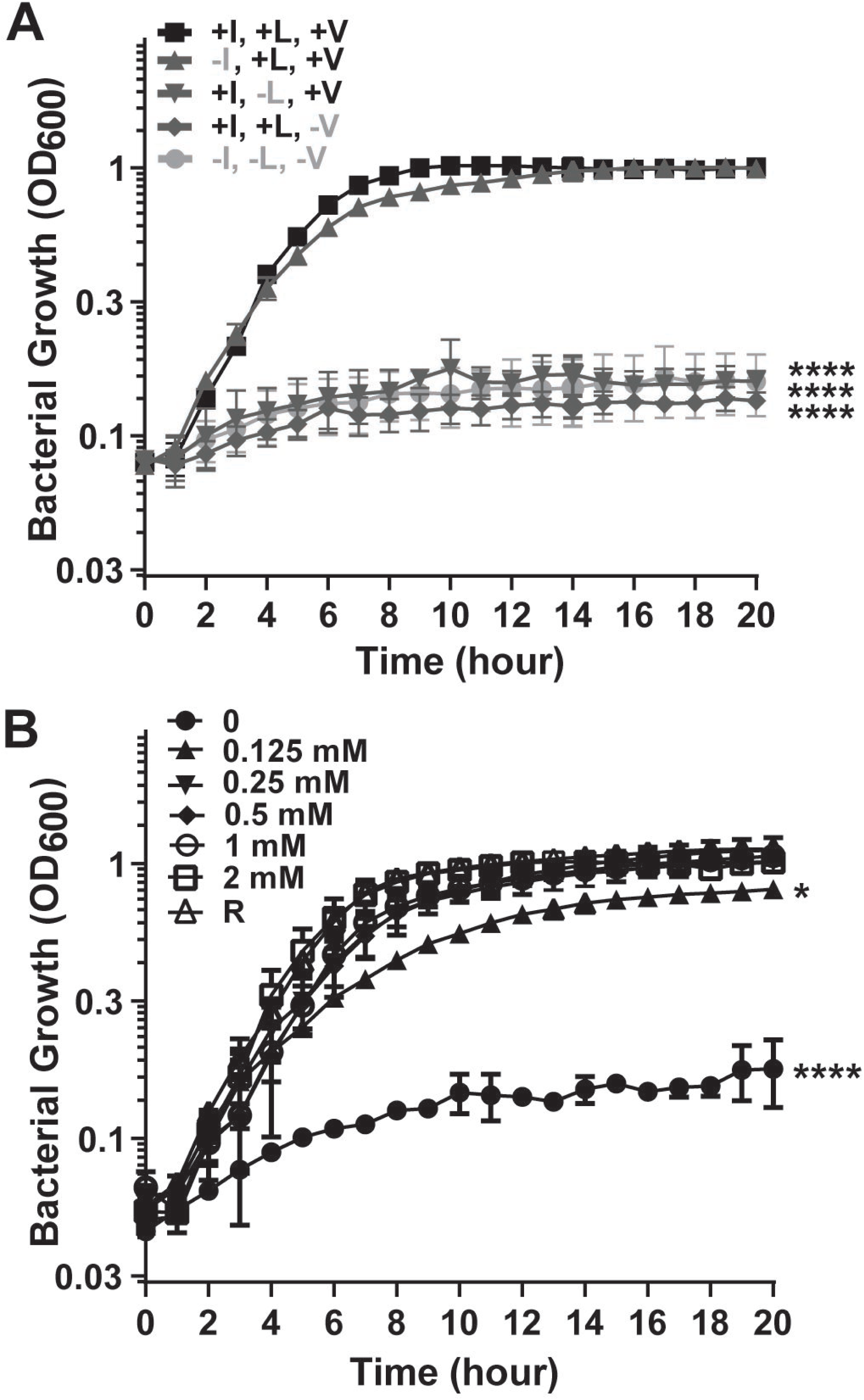
Requirement of BCAAs for *B. anthracis* ANR-1 growth in toxin-inducing conditions. (A) Growth in R medium (1.75 mM isoleucine, 1.5 mM leucine, and 1.35 mM valine) and R medium missing one or more BCAAs as indicated. (B) Growth in R medium with altered concentrations of BCAAs. Each of the three BCAAs were present at the concentrations indicated. Data are the mean of three biological replicates with error bars representing standard deviation. Data were compared with the growth in R medium and analyzed using one-way Analysis of variance (ANOVA) followed by Dunnett’s multiple comparisons analysis. Asterisks indicate P-values. (* =P<0.05, **** =P<0.0001).

To explore BCAA concentrations required for growth in toxin-inducing culture conditions, we cultured *B. anthracis* in R medium and R medium with altered concentrations of BCAAs (Fig. 1B). Again, the absence of all BCAAs resulted in poor growth. Growth was restored when all BCAAs were present at 0.125 mM, and BCAA concentrations of at least 0.25 mM resulted in growth rates and yields comparable to those obtained in R medium. The importance of BCAAs for optimal growth of *B. anthracis* in toxin-inducing conditions prompted us to examine the genome for predicted BCAA biosynthesis and transport genes.

### Expression of BCAA biosynthesis genes of *B. anthracis*

The interconnected pathways for isoleucine, leucine, and valine biosynthesis are highly conserved in bacteria (18, 23). Our analysis of the annotated *B. anthracis* genome revealed two loci, that we designated *ilv1* and *ilv2*, that together contain all genes required for the biosynthesis of BCAAs (12, 13). Some BCAA biosynthesis genes have alleles in both loci, while others exist in a single copy, as depicted in figure 2A. Both loci carry copies of *ilvE*, *ilvB*, and *ilvC*. Enzymes encoded by these genes are required for synthesis of all three BCAAs. By contrast, *ilvD* encoding another enzyme needed for synthesis of all BCAAs is present in single copy on *ilv2*. The *ilv2* locus also contains *ilvA*, encoding an enzyme specific for isoleucine biosynthesis. Genes specific for leucine biosynthesis, *leuA*, *leuB*, *leuC* and *leuD*, are clustered together in the *ilv1* locus. Notably, while the sequences of *B. anthracis* BCAA biosynthesis genes are indicative of proteins that are highly homologous to the well-characterized enzymes of *B. subtilis*, the gene arrangement in *B. anthracis* differs from that found in *B. subtilis*. The BCAA biosynthesis genes of *B. subtilis* are found in three loci: the *ilv-leu* operon, the *ilvA* gene, and the *ilvD* gene, and all of the genes are present in single copy (46).

**Figure 2:**
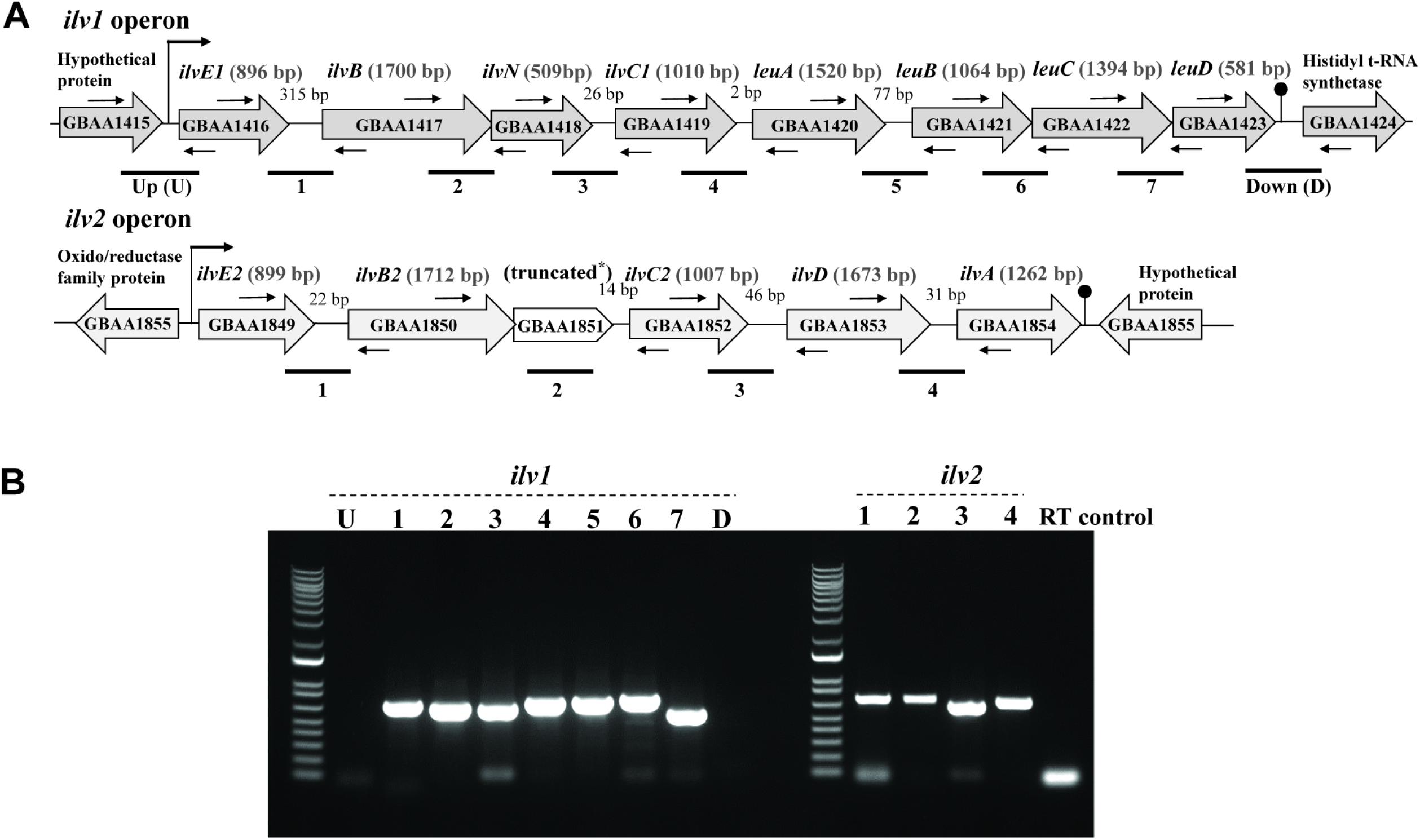
Organization and expression of *ilv* loci in *B. anthracis*. (A) Schematic representation of the *ilv* loci. Two operons designated as operon *ilv*1 and operon *ilv*2 are shown. Genes are indicated as open arrows with the corresponding annotations (https://www.ncbi.nlm.nih.gov/nucleotide/AE017334). Gene sizes (bp) are shown in parentheses. Truncated* is the truncation of a gene predicted to encode the small subunit of acetolactate synthase III. Sizes of intergenic spaces (bp) are as indicated. Predicted transcription start and termination sites are denoted by thick arrows and hairpins, respectively. Thin horizontal arrows represent primer pairs for PCR. Small thin arrows above and below genes indicate the approximate positions of primers used for RT-PCR. Horizontal lines below the operons correspond to the RT-PCR products shown in B. (B) Co-transcription of *ilv* loci. Ethidium bromide-stained agarose gels showing the RT-PCR products obtained using primers shown in A. Lane designations correspond to anticipated products shown in A. The RT control reaction contained RNA as a template. Primers directly upstream (U) and downstream (D) of the *ilv1* operon were used as negative controls for co-transcription with genes flanking the *ilv1* operon in the same DNA strand.

We performed RT-PCR experiments to assess expression of the BCAA biosynthesis genes. Using RNA from cultures grown in R medium with 0.25 mM BCAAs and primers corresponding to intergenic and flanking regions, we detected RT-PCR products indicating co-transcription of all genes within each locus (Fig. 2B). Thus, *ilv1* and *ilv2* represent operons.

Transcription of the BCAA biosynthesis operons was also detected in RNA from cultures grown in complete R medium (data not shown).

### Predicted BCAA transport genes of *B. anthracis*

Despite expression of BCAA biosynthesis genes, our experiments showed apparent BCAA auxotrophy during culture in defined media (Fig. 1). The importance of exogenous BCAAs for culture of *B. anthracis* prompted us to examine the *B. anthracis* genome for predicted BCAA transporter genes. Our bioinformatic analyses revealed that *B. anthracis* carries an unusually large number of predicted BCAA transporter genes compared to other Gram-positive bacteria for which BCAA transport has been studied. *B. anthracis* harbors six genes, *brnQ1* (GBAA_690), *brnQ2* (GBAA_0802), *brnQ3* (GBAA_1459), *brnQ4* (GBAA_2063), *brnQ5* (GBAA_3142), and *brnQ6* (GBAA_4790), predicted to encode BCAA transport system carrier II proteins. These proteins are LIVCS (Leucine, isoleucine and valine cationic symporter) family proteins which function by a Na^+^ or H^+^ symport mechanism (47). They typically contain 12 transmembrane helices and are members of the major facilitator superfamily (MFS) (18, 48). The genome also contains five genes, GBAA_1931, GBAA_1933, GBAA_1934, GBAA_1935, and GBAA_1936, predicted to be ABC transporters for BCAAs, as well as another gene, GBAA_0818, that is similar to *bcaP*, an amino acid permease gene that is found in many Gram-positive bacteria (16, 29, 31, 49). By contrast, the pathogens *S. aureus* and *S. pneumoniae* harbor four and one known or predicted BCAA transporters, respectively (16, 21, 22). The non-pathogen *B. subtilis* harbors three BCAA transporter genes (29). *C. glutamicum*, a bacterium used for industrial synthesis of amino acids, only carries one BCAA transport gene (30) and *L. lactis*, used in the production of fermented milk products, harbors two BCAA transport genes (31). The relatively large number of apparent BCAA transporter genes of *B. anthracis* supports the importance of BCAA transport in this bacterium.

### Characterization of *brnQ*-null mutants

The presence of multiple genes predicted to encode BCAA transporters coupled with previous reports of controlled expression of the *B. anthracis brnQ* genes in response to regulators and signals associated with virulence (4, 15, 33, 34) spurred us to investigate BrnQ function. All six BrnQ paralogs are similar in length [BrnQ1: 433 amino acids (aa), BrnQ2: 438 aa, BrnQ3: 451 aa, BrnQ4: 445aa, BrnQ5: 450 aa, BrnQ6: 441 aa] and they share at least 42% sequence identity and 62% sequence similarity. Each protein is predicted to have 12 transmembrane domains except for BrnQ4 which has 11 such domains (TMPred program: https://embnet.vital-it.ch/software/TMPRED_form.html) (50). We created individual *brnQ*-null mutants and compared their growth to the parent strain when cultured in R medium under toxin-inducing conditions. All six mutants showed growth rates comparable to the parent, revealing that no single *brnQ* gene is essential for *B. anthracis* growth in culture conditions in which BCAA transport is required (Fig 3).

**Figure 3:**
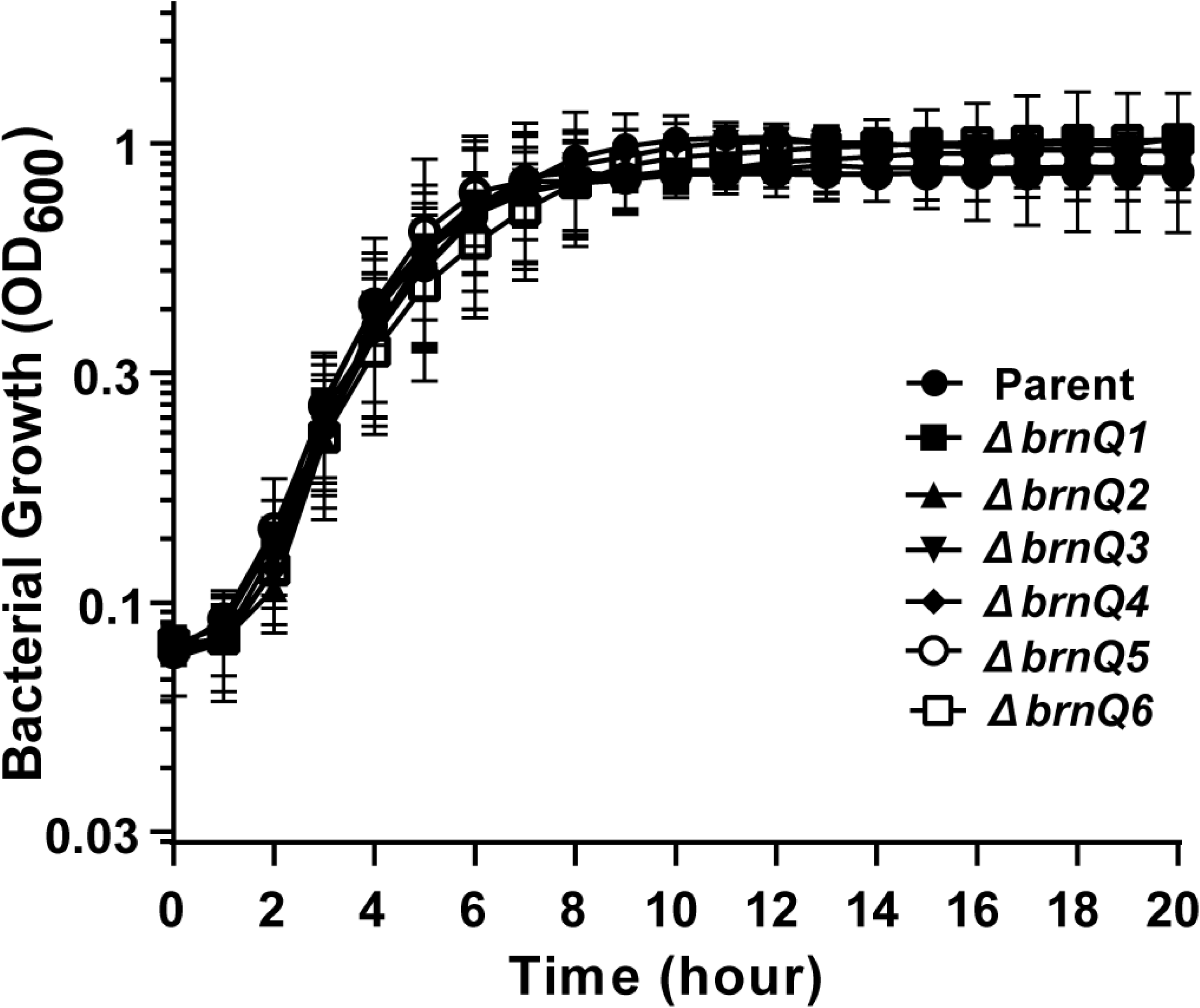
Growth of single *brnQ* transporter null-mutants in R medium. Each data-point represents the average for three independent experiments +/- standard deviation. Data were analyzed using one-way Analysis of variance (ANOVA) followed by Dunnett’s multiple comparisons and compared with the parent strain.

We performed BCAA transport assays to assess the ability of the parent strain, individual *brnQ*-null mutants, and complemented mutants to uptake radiolabeled isoleucine, leucine, and valine. For complementation of the null mutations, the appropriate genes were cloned under control of a xylose-inducible promoter and expression was induced with 1% xylose. Although the assays were performed using native *brnQ* genes in the complemented mutants, we verified the induction system by cloning recombinant *brnQ* genes carrying a FLAG tag at the 3’ terminus and protein expression was confirmed by immunoblotting with anti-FLAG antibody.

Results of the BCAA transport assays are shown in figure 4. A strong phenotype was associated with *brnQ4*. Isoleucine and valine uptake were reduced substantially in the *brnQ4*- null mutant compared to the parent strain, and the phenotype was partially complemented by expression of *brnQ4 in trans*. Deletion of *brnQ4* did not affect leucine uptake. These data suggest that BrnQ4 is a major transporter for isoleucine and valine under these growth conditions. Deletion of *brnQ3* also resulted in a statistically significant decrease in isoleucine and valine uptake, and expression of *brnQ3 in trans* restored the mutant to the parent phenotype. Interestingly, the *brnQ3*-null mutant exhibited increased leucine uptake, a phenotype that was not complemented by *brnQ3 in trans*. Deletion of *brnQ1* resulted in a small but statistically significant decrease in isoleucine and valine uptake, however complementation by expression of *brnQ1 in trans* was not achieved. Individual deletions of the other *brnQ* genes did not result in statistically significant changes in BCAA uptake.

**Figure 4:**
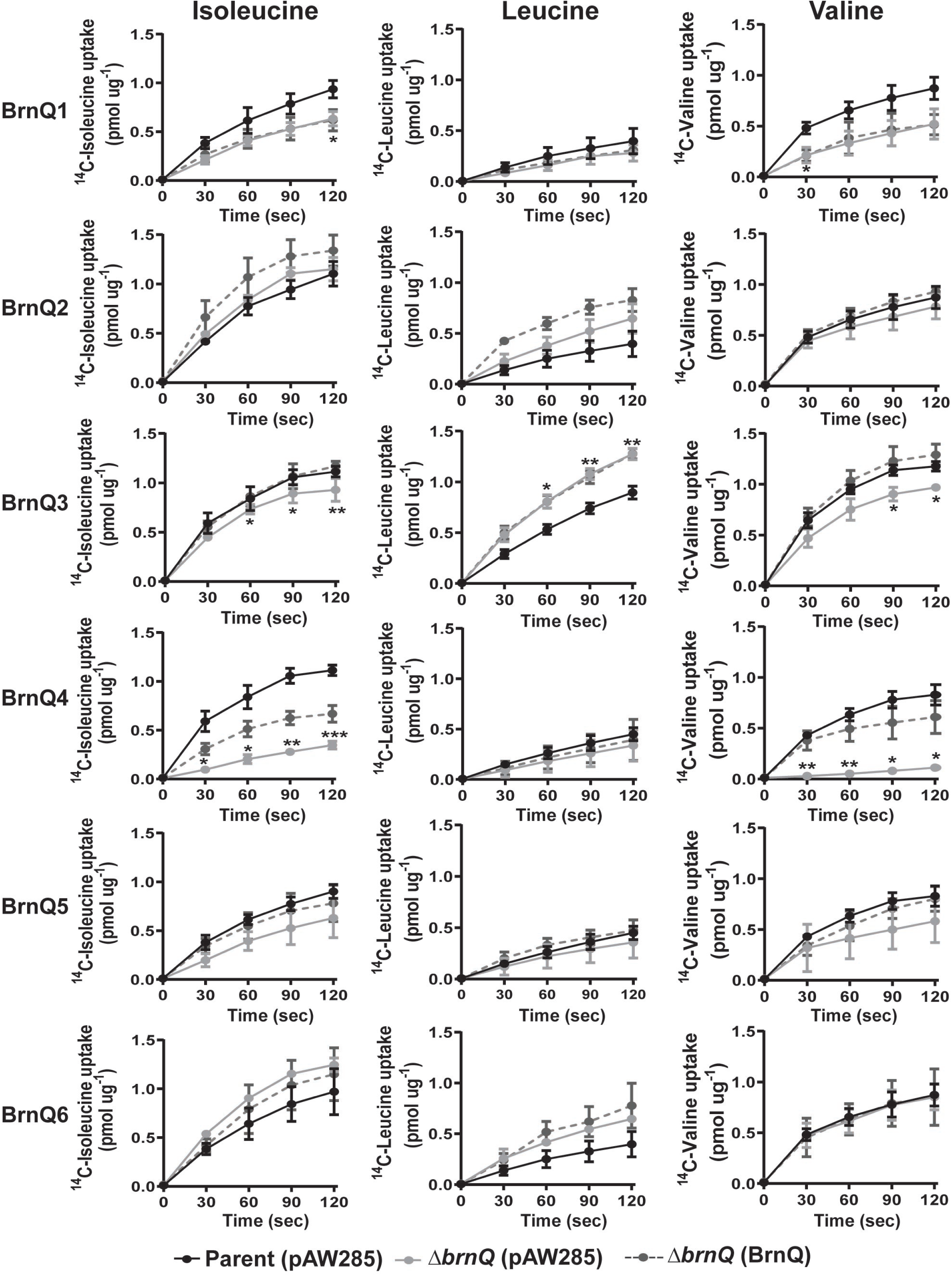
BCAA transport by single *brnQ*-null mutants. Transport of ^14^C-labeled isoleucine, ^14^C-labeled leucine, and ^14^C-labeled valine was assessed for the parent strain containing empty vector pAW285 (black solid line), individual *brnQ* mutant with pAW285 (grey solid line), and individual *brnQ* mutant complemented with corresponding gene (grey broken line). Data represent the means from three biological replicates +/- standard deviations. Data were analyzed using two-way Analysis of Variance (ANOVA) with repeated measures followed by Bonferroni’s multiple comparisons analysis. Comparisons of parent and single-deletion mutants are shown with asterisks representing P-values. (* =P<0.05, ** =P<0.01, *** =P<0.001).

Overall, our assessment of growth and BCAA uptake by the single *brnQ*-null mutants suggests that multiple BrnQ transporters with varying levels of functional redundancy and specificity are involved in BCAA uptake.

### Specificity of BrnQ transporters

To explore specificity of BrnQ transporters, we individually expressed *brnQ* genes *in trans* in a mutant deleted for multiple *brnQ* genes that was also deficient for BCAA biosynthesis. We reasoned that assessment of transporter activity might be enhanced in a mutant unable to synthesize BCAAs. To construct the mutant, we first deleted *ilvD*. The *ilvD* gene encodes dihydroxy acid dehydratase, an enzyme essential for biosynthesis of all three BCAAs. The growth rate of the *ilvD*-null mutant in R medium was reduced compared to that of the parent strain, but after 12 h the densities of the two cultures were comparable (Fig. 5A). This result agrees with our previous data (Fig. 1 and Fig. 2B) suggesting that while BCAA transport is sufficient to achieve optimal growth, BCAA biosynthesis is still active in media containing BCAAs.

**Figure 5:**
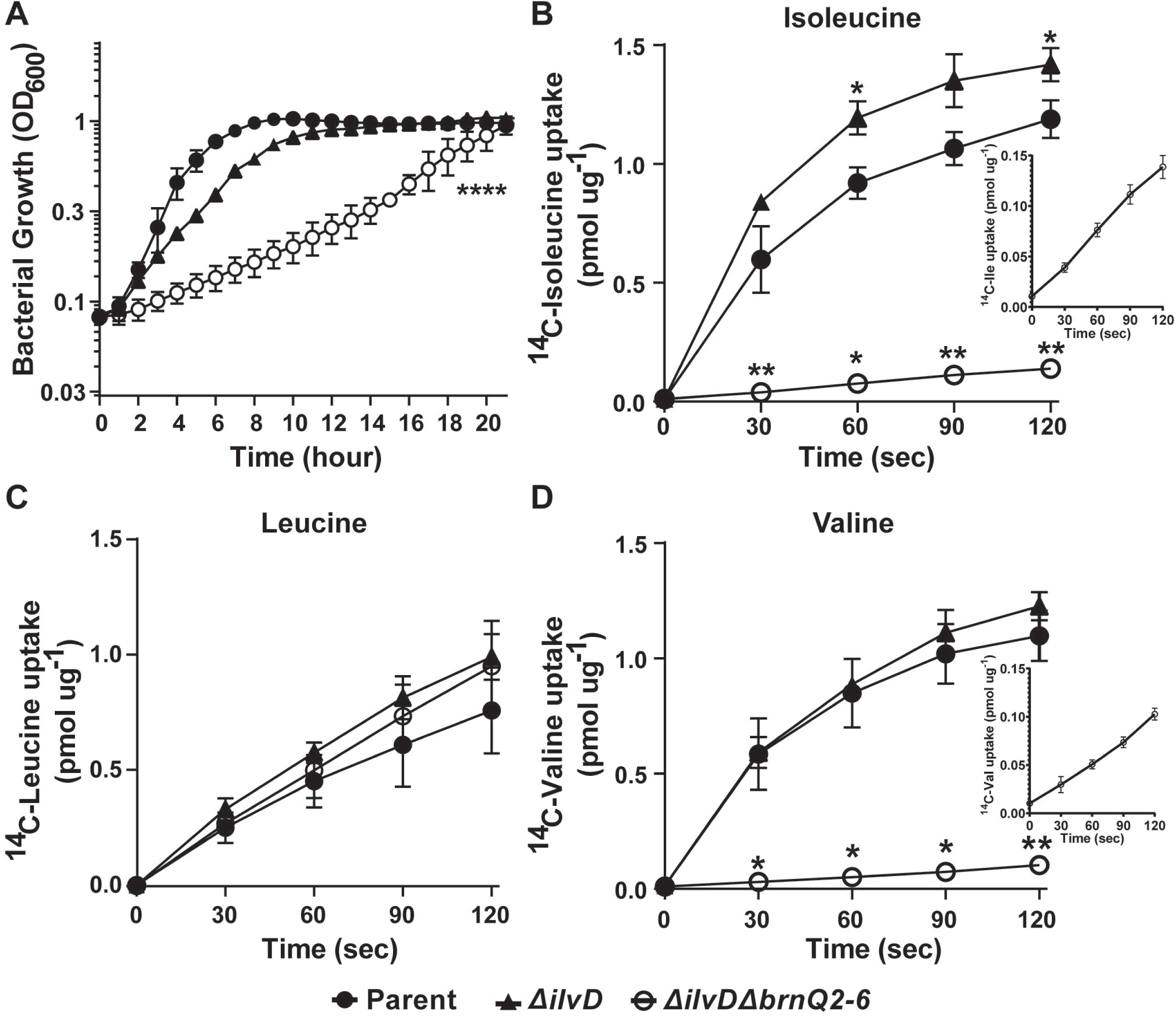
Growth and BCAA transport by the *ΔilvD* and *ΔilvDΔbrnQ2-6* mutants. (A) Growth of *ΔilvD and ΔilvDΔbrnQ2-6* mutant in R medium. Data are represented as the mean from three independent experiments. Growth was compared to the respective growth of parent. Error bars represent standard deviations. Data were analyzed using one-way Analysis of Variance (ANOVA) followed by Dunnett’s multiple comparisons analysis. Asterisks indicate P-values. **** =P<0.0001). (B-D) BCAA uptake. Uptake of (B) ^14^C-labeled isoleucine, (C) ^14^C-labeled leucine, and (D) ^14^C-labeled valine was assessed for the parent (closed circle), *ΔilvD* (triangle), and *ΔilvDΔbrnQ2-6* (open circle) strains. Uptake of isoleucine and valine by *ΔilvDΔbrnQ2-6* mutant is shown in the inset (B and D respectively). Data are the means from three biological replicates +/- standard deviations. Data were analyzed using two-way Analysis of Variance (ANOVA) with repeated measures followed by Bonferroni’s multiple comparisons analysis. Comparison of parent with the *ΔilvD* mutant, and parent with the *ΔilvDΔbrnQ2-6* mutant was assessed and shown in asterisks indicating P values (* =P<0.05, ** = P<0.01).

Multiple attempts to create an *ilvD*-null mutant deleted for all 6 *brnQ* genes were unsuccessful, suggesting that the presence of at least one *brnQ* gene is essential for BCAA transport. Successive deletion of *brnQ2, brnQ3, brnQ4, brnQ5,* and *brnQ6* in the *ilvD*-null background resulted in a mutant, *ΔilvDΔbrnQ2-6*, that exhibited a severe growth defect in R medium (Fig. 5A), likely due to significantly reduced BCAA transport. BCAA uptake by the *ΔilvDΔbrnQ2-6* mutant was compared to the parent strain and the *ilvD*-null mutant (Fig. 5B-D). As expected, BCAA uptake was enhanced in the *ilvD* mutant compared to the parent, with a statistically significant increase in isoleucine uptake and small but reproducible increases in leucine and valine uptake. The *ΔilvDΔbrnQ2-6* mutant showed greatly reduced isoleucine and valine uptake (Fig. 5B and 5D), suggesting that one or more of the deleted *brnQ* genes encode major transporters of these BCAAs and that the reduced growth rate of the *ΔilvDΔbrnQ2-6* mutant in R medium (Fig. 5A) is related to poor uptake of isoleucine and valine by less active transporters. The *ΔilvDΔbrnQ2-6* mutant was unaffected for leucine uptake (Fig. 5C), indicating that *brnQ1* and/or unidentified transporters are associated with leucine transport.

Next, we tested for complementation of the *ΔilvDΔbrnQ2-6* mutant phenotype by individual *brnQ* genes. We assessed isoleucine and valine transport associated with expression of *brnQ*2, *brnQ3*, *brnQ4*, *brnQ5*, and *brnQ6 in trans*. Expression of *brnQ3*, *brnQ4*, and *brnQ5* partially restored isoleucine and valine uptake by the *ΔilvDbrnQ2-6* mutant, while expression of *brnQ2* and *brnQ6* did not complement the uptake deficiency of the *ΔilvDΔbrnQ2-6* mutant (Fig. 6). Taken together, our results indicate that BrnQ3, BrnQ4 and BrnQ5 can act as isoleucine and valine transporters.

**Figure 6:**
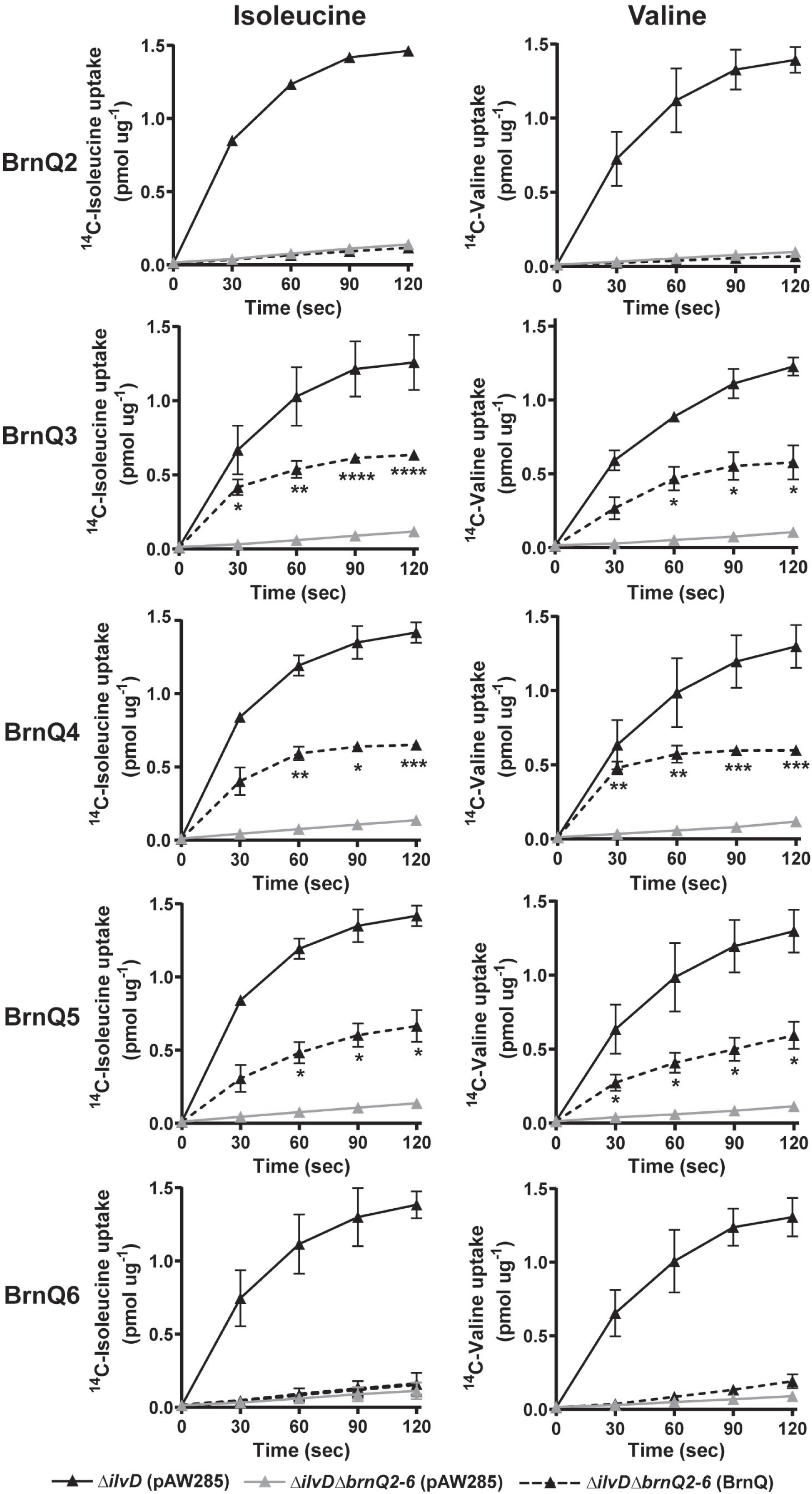
Isoleucine and Valine transport associated with expression of specific BrnQs. Transport of ^14^C-labeled isoleucine and valine was assessed for the *ΔilvD* mutant containing empty pAW285 vector (black solid line), the *ΔilvDΔbrnQ2-6* mutant containing empty pAW285 vector (grey solid line), and the *ΔilvDΔbrnQ2-6 strain* complemented with individual *brnQ* genes (black broken line). Values represent the mean from three independent experiments +/- standard deviations. Data were analyzed using two-way Analysis of Variance (ANOVA) with repeated measures followed by Bonferroni’s multiple comparisons analysis. Each complemented BrnQ+ mutant was compared to the *ΔilvDΔbrnQ2-6* mutant and shown in asterisks indicating P values (* =P<0.05, ** = <0.01, *** =P<0.001, **** =P<0.0001).

To compare the activities of BrnQ3, BrnQ4, and BrnQ5, we measured the initial velocities of BCAA uptake in increasing concentrations of isoleucine and valine by the *ΔilvDΔbrnQ2-6* mutant expressing individual *brnQ3, brnQ4,* or *brnQ5* genes (Fig. 7). Apparent K_m_ and V_max_ values associated with each transporter are shown in Table 2. The data indicate that the three transporters have comparable affinities for isoleucine and valine under our experimental conditions. The highest initial velocity was associated with BrnQ4, suggesting that BrnQ4 is the major determinant for isoleucine and valine transport, which is in agreement with the strong phenotype of the *brnQ4*-null mutant in our BCAA transport assays (Fig. 4). Altogether, our results establish roles for BrnQ3, BrnQ4 and BrnQ5 in BCAA transport by *B. anthracis* cultured in toxin-inducing conditions.

**Figure 7:**
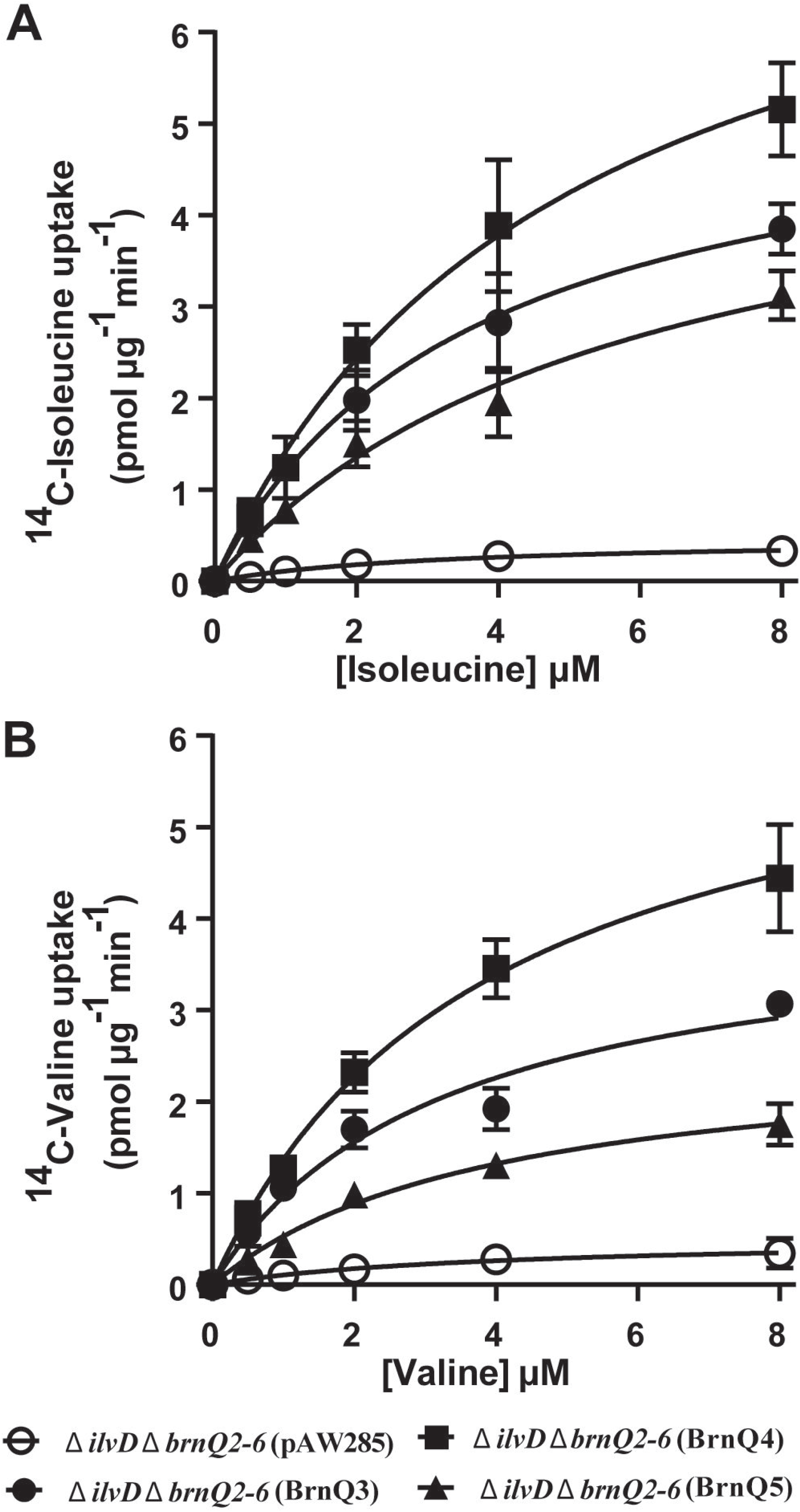
Kinetics of isoleucine and valine transport associated with BrnQ3, BrnQ4, and BrnQ5. Isoleucine (A) and valine (B) uptake kinetics are assessed for *ΔilvDΔbrnQ2-6* with empty vector pAW285 (open circles), *ΔilvDΔbrnQ2-6 strain* with BrnQ3 (closed circles), *ΔilvDΔbrnQ2-6 strain* with BrnQ4 (closed squares), and *ΔilvDΔbrnQ2-6 strain* with BrnQ5 (closed triangles). Data shown represent the mean of three independent experiments. Error bars represent +/- standard deviations.

### Role of BCAA transport and synthesis in virulence

To assess the significance of BrnQ3, BrnQ4, and BrnQ5 during *B. anthracis* infection, we tested *brnQ*-null mutants for virulence and tissue burden in a murine model for systemic anthrax. Groups of 7-8 weeks-old complement-deficient female A/J mice were infected intravenously with ∼10^5^ CFUs of the parent strain or single *brnQ*-null mutants. Mice were monitored for up to 11 days. All parent-infected mice succumbed to anthrax disease within eight days of infection (Fig. 8A). The time to death for mice infected with *brnQ4*- and *brnQ5*-null mutants was not significantly different than the time to death for mice infected with the parent strain. However, the *brnQ3*-null mutant was highly attenuated in this animal model. Nine of ten mice infected with this mutant survived infection and exhibited no signs of disease.

**Figure 8:**
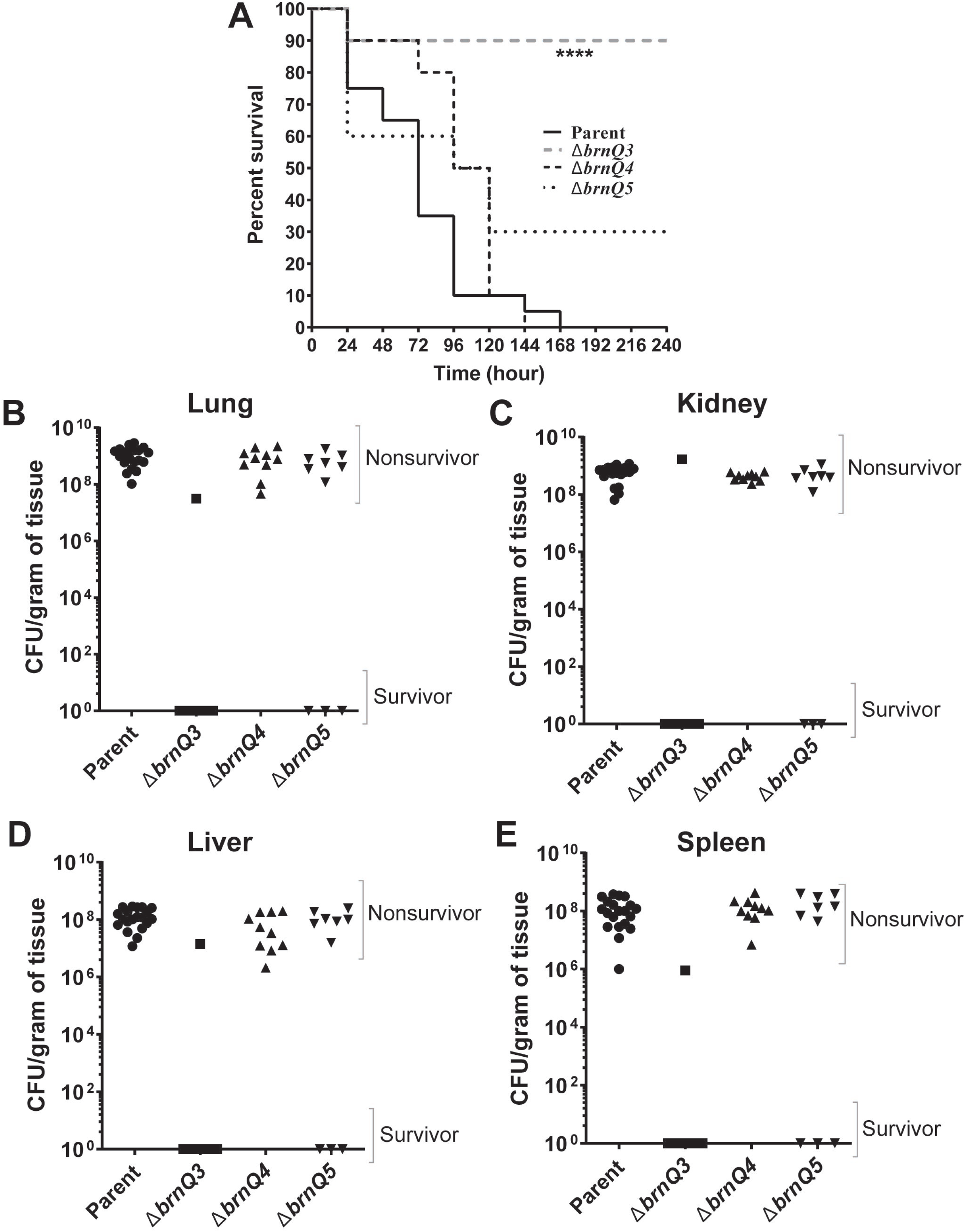
Virulence of BCAA transporter mutants. 7-8 weeks old female A/J mice were infected with ∼10^5^ CFU of ANR-1 parent strain (n=20), *ΔbrnQ3* mutant (n=10), *ΔbrnQ4* mutant (n=10) or *ΔbrnQ5* mutant (n=10) via tail-vein injection. Mice were monitored for 11 consecutive days. Organs were collected from dead and surviving mice, homogenized, and plated for determination of CFU. (A) Kaplan-Meier survival curves are shown for parent (black solid line), *ΔbrnQ3* mutant (grey broken line), *ΔbrnQ4* mutant (black broken line), and *ΔbrnQ5* mutant (black dotted line). Statistical significance was analyzed using the log-rank (Mantel-Cox) test and compared with the parent. P-value is indicated by asterisk (**** = P<0.0001). (B-E) CFUs in lungs (B), kidneys (C), livers (D), and spleens (E) from non-survivors and survivors were determined for parents (circles), *ΔbrnQ3* mutant (squares), *ΔbrnQ4* mutant (triangles), and *ΔbrnQ5* mutant (inverted triangles). No detectable CFUs were found in organs of survivors. Individual data points are shown. Data for each mutant were compared to parent. Mann-Whitney unpaired Student’s t-test was used to determine the significance.

We collected the lungs, kidneys, livers, and spleens from surviving and diseased mice to determine bacterial burdens in tissues. For all mice that succumbed to infection, comparable numbers of CFUs were found in tissues infected with the parent and mutants (Fig. 8B-E). Approximately 10^9^ CFU per gram of tissue was found in the lung and kidney, and approximately 10^8^ CFU per gram of tissue was found in the liver and spleen. Thus, for diseased mice, the absence of a single isoleucine/valine transporter did not affect numbers of the bacterium in various tissues. No detectable CFU were found in mice that survived infection.

Overall, these data show that BrnQ3 is required for full virulence even in the presence of other isoleucine/valine transporters, BrnQ4 and BrnQ5. The results indicate that despite the ability of BrnQ3, BrnQ4, and BrnQ5 to transport isoleucine and valine in cultured cells, these transporters do not have redundant activity during infection.

Our data showed that deletion of *ilvD* has a minimal effect on *B. anthracis* growth in culture when all transporter genes are present. To determine if BCAA biosynthesis is important during infection, we also tested the *ilvD*-null mutant for virulence. Surprisingly, the *ilvD*-null mutant was attenuated (Fig. 9A). Eighty percent of the mice injected with the *ilvD*-null mutant did not exhibit symptoms, survived infection, and had no recoverable CFU in their tissues. For *ilvD*-null infected mice that succumbed to anthrax disease, the numbers of CFU in tissue from the lung, kidney, liver and spleen were similar to those found in mice infected with the parent strain (Fig. 9B-E). These results suggest that while *ilvD* is not essential for growth in culture medium containing BCAAs, *ilvD* plays a role in virulence in the murine model.

**Figure 9:**
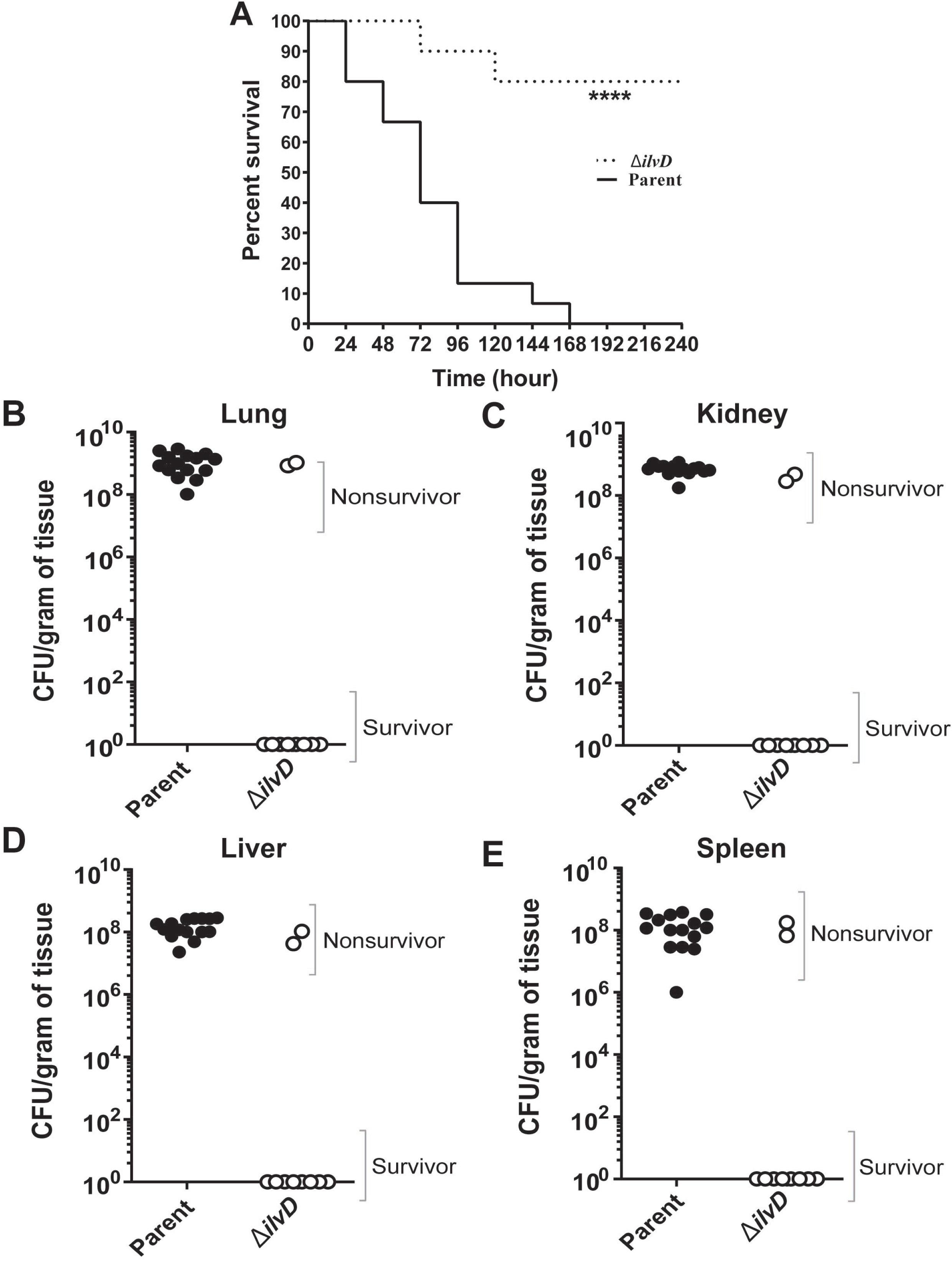
Virulence of BCAA biosynthesis mutant. Mice were infected as described in Fig. 8. Parent strain (n=15), *ilvD*-null mutant (n=10). (A) Kaplan-Meier survival curves are represented by a solid line for the parent and by a dotted line for the *ΔilvD* mutant. Statistical significance was analyzed by the log-rank (Mantel-Cox) test and P-value is denoted by asterisks (****=P<0.0001). (B-E) CFU values in collected lungs (B), kidneys (C), livers (D), and spleens (E) of non-survivors and survivors were represented for parent (closed circles) and *ilvD*-null mutant (open circles). Significance of the differences was analyzed by Mann-Whitney unpaired Student’s t-test.

### BCAA levels in mouse tissues

Considering the attenuated virulence of a mutant deficient in BCAA biosynthesis and a mutant lacking the isoleucine/valine transporter BrnQ3, we sought to determine the relative abundance of BCAAs in mouse tissues. BCAA availability in mammalian tissues is largely undefined (18). Tissues from three uninfected mice were harvested, amino acids were extracted, and samples were analyzed by LC-MS. The relative abundance of each BCAA was normalized with tissue weight (Table 3, Figure S1.) Of the four organs tested, the spleen contained the greatest concentration of all three BCAAs. For each organ, valine was the most abundant BCAA, while isoleucine was the least abundant BCAA. Overall, only approximately 2- to 3-fold differences were noted between organs, suggesting somewhat comparable BCAA availability in these niches.

### Influence of BCAAs on AtxA activity

AtxA is a critical positive regulator of the anthrax toxin genes and the *B. anthracis* capsule biosynthesis operon (35, 37, 51). In our murine model for anthrax, death of the animal is highly dependent upon synthesis of the anthrax toxin proteins, and an *atxA-*null mutant, which is toxin-deficient, is avirulent (36, 38). Our more recent investigations show that in addition to control of toxin and capsule genes, AtxA negatively affects expression of BCAA transport- and synthesis-related genes (15), most likely via an indirect mechanism dependent upon a small regulatory RNA, XrrA (4). Transcriptional profiling data suggest that AtxA activates transcription of XrrA (15), and XrrA represses the BCAA genes (4). To address potential relationships between BCAAs and AtxA that could be associated with attenuated virulence of the *ilvD*- and *brnQ3*-null mutants, we assessed *atxA* expression and AtxA activity during culture of *B. anthracis* in different levels of BCAAs.

To measure *atxA* promoter activity, we performed β-galactosidase assays using ANR-1 (pUTE839), which carries an *atxA* promoter-*lacZ* fusion (Table 1). As shown in Fig 10A, no significant differences in β-galactosidase activity were detected in cultures grown in R medium with BCAA concentrations ranging from 0.25 mM to 4 mM.

**Figure 10:**
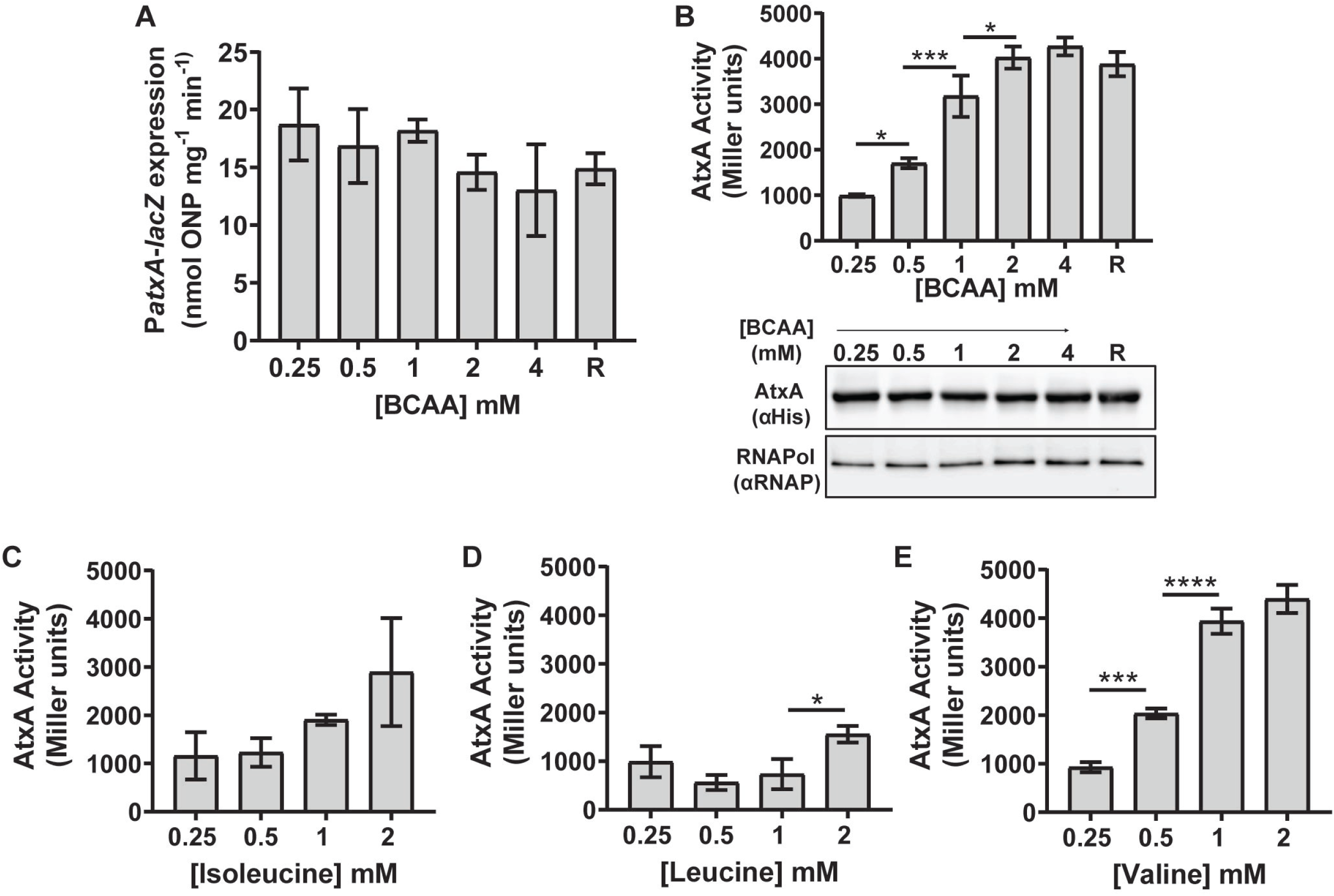
Effect of BCAAs on *atxA* promoter activity, AtxA function, and AtxA protein level. (A) *atxA* promoter activity. *atxA* promoter activity of a reporter strain carrying a P*atxA*- *lacZ* transcriptional fusion was measured using a β-galactosidase assay following growth in varying BCAA concentrations. (B) AtxA protein activity *in vivo*. An *atxA*-null strain (UT376) carrying P*lef-lacZ* (a reporter for AtxA activity) and containing IPTG-inducible His-tagged *atxA* allele was induced during growth in R medium of varying BCAA concentrations. β- galactosidase activity (upper panel) and AtxA protein levels were measured (lower panel).. Steady state levels of AtxA and RNA polymerase β-subunit were detected in cell lysates via immunoblotting using α-His antibody and α-RNA Pol β-antibody respectively. The experiment was performed three times and a representative image is shown. (C-E) Effects of isoleucine (C), leucine (D), and valine (E) on AtxA activity. β-galactosidase assays were performed as described in (B). The concentration of one BCAA (as indicated) was altered while keeping the concentrations of the other BCAAs constant at 0.25 mM to support optimal growth. Each bar represents the mean of three biological replicates. Error bars represent +/- standard deviation. One-way ANOVA followed by Tukey’s multiple comparisons tests were performed to analyze the data. Asterisks indicate P-values. (* =P<0.05, *** = <0.001, **** =P<0.0001).

**Table 1:** *B. anthracis* strains and plasmids

| Strains | Relevant Characteristics | Source |
| --- | --- | --- |
| ANR-1 | *B. anthracis* parent strain, pXO1<sup>+</sup>, pXO2<sup>-</sup> | (72) |
| UT376 | ANR-1-derivative, *atxA*-null, *lef* promoter- *lacZ* fusion, (*Plef-lacZ*) at native *lef* locus | (39) |
| UT441 | ANR-1-derivative, $\Delta brnQ3$ | This work |
| UT469 | ANR-1-derivative, $\Delta ilvD$ | This work |
| UT472 | ANR-1-derivative, $\Delta brnQ2$ | This work |
| UT475 | ANR-1-derivative, $\Delta brnQ5$ | This work |
| UT476 | ANR-1-derivative, $\Delta brnQ6$ | This work |
| UT478 | ANR-1-derivative, $\Delta brnQ4$ | This work |
| UT481 | ANR-1-derivative, $\Delta brnQ1$ | This work |
| UT482 | ANR-1-derivative, $\Delta ilvD brnQ2 brnQ3 brnQ4 brnQ5 brnQ6$ ($\Delta ilvD \Delta brnQ2-6$) | This work |
| Plasmids |  |  |
| pHY304 | Temperature-sensitive vector used for deletion of indicated loci by homologous recombination; Erm<sup>R</sup> | (73) |
| pAW285 | Xylose inducible expression vector, Cm<sup>R</sup> | (70) |
| pUTE839 | pHT304-18z-derived *atxA* promoter-lacZ fusion vector containing sequence from -770 to +99 | (38) |
| pUTE991 | pUTE657-derived expression vector for AtxA-His<sub>6</sub> (6 x His-epitope on the C-terminus of AtxA) | (39) |
| pUTE1165 | pAW285-derived expression vector for BrnQ3-FLAG (FLAG-tag on the C-terminus of BrnQ3) | This work |
| pUTE1167 | pAW285-derived expression vector for BrnQ6-FLAG (FLAG-tag on the C-terminus of BrnQ6) | This work |
| pUTE1191 | pAW285-derived expression vector for BrnQ2 | This work |
| pUTE1192 | pAW285-derived expression vector for BrnQ3 | This work |
| pUTE1193 | pAW285-derived expression vector for BrnQ6 | This work |
| pUTE1206 | pAW285-derived expression vector for BrnQ1 | This work |
| pUTE1207 | pAW285-derived expression vector for BrnQ4 | This work |
| pUTE1208 | pAW285-derived expression vector for BrnQ5 | This work |
| pUTE1209 | pAW285-derived expression vector for BrnQ1-FLAG (FLAG-tag on the C-terminus of BrnQ1) | This work |
| pUTE1210 | pAW285-derived expression vector for BrnQ2-FLAG (FLAG-tag on the C-terminus of BrnQ2) | This work |
| pUTE1211 | pAW285-derived expression vector for BrnQ4-FLAG (FLAG-tag on the C-terminus of BrnQ4) | This work |
| pUTE1212 | pAW285-derived expression vector for BrnQ5-FLAG (FLAG-tag on the C-terminus of BrnQ5) | This work |

**Table 2:** Apparent K_m_ and V_max_ values for BrnQ3, BrnQ4 and BrnQ5 transporters.

| | Apparent $K_m$<br>($\mu M$) | | Apparent $V_{max}$<br>($pmol\ \mu g^{-1}\ min^{-1}$) | |
| --- | --- | --- | --- | --- |
|  | Isoleucine | Valine | Isoleucine | Valine |
| **BrnQ3** | $3.6 \pm 1.6$ | $3.3 \pm 1.8$ | $5.5 \pm 1.2$ | $4.1 \pm 0.9$ |
| **BrnQ4** | $4.9 \pm 2.9$ | $3.9 \pm 1.6$ | $8.5 \pm 2.5$ | $6.7 \pm 1.3$ |
| **BrnQ5** | $5.8 \pm 3.7$ | $4 \pm 2.5$ | $5.3 \pm 2.3$ | $2.6 \pm 0.8$ |

**Table 3:** BCAA levels in several mouse tissues

| | **Lung**<br>($nmol\ mg^{-1}\ tissue$) | **Liver**<br>($nmol\ mg^{-1}\ tissue$) | **Spleen**<br>($nmol\ mg^{-1}\ tissue$) | **Kidney**<br>($nmol\ mg^{-1}\ tissue$) |
| --- | --- | --- | --- | --- |
| **Isoleucine** | 0.08-0.13 | 0.13-0.18 | 0.19-0.23 | 0.12-0.14 |
| **Leucine** | 0.15-0.2 | 0.22-0.28 | 0.39-0.41 | 0.21-0.22 |
| **Valine** | 0.2-0.32 | 0.26-0.31 | 0.46-0.54 | 0.27-0.30 |

We quantified AtxA activity using reporter strain UT376, an *atxA*-null mutant harboring an IPTG-inducible His-tagged *atxA* allele *in trans*. This strain also contains a transcriptional fusion in which the promoter of an AtxA-regulated gene, *lef*, is fused to a promoter-less *lacZ*, at the native *lef* locus. As shown in figure 10B (upper panel), BCAAs enhanced AtxA activity in a dose-dependent manner, with maximum activity at 2 mM BCAAs, which is comparable to the BCAA concentration in R medium. Results of western blotting experiments indicated that the AtxA protein levels were similar in cultures containing different BCAA concentrations (Fig. 10B, lower panel).

We repeated the AtxA activity experiments, varying the level of either isoleucine, leucine or valine, while keeping the other two BCAAs at the minimum concentration required to support optimal growth (0.25 mM). Valine exhibited a statistically significant dose-dependent effect on AtxA activity (Fig. 10E), while the effects of isoleucine and leucine were variable (Figs. 10C-D). These data show that BCAAs, and specifically valine, increase AtxA activity in a dose-dependent manner, linking BCAAs with virulence factor expression.

## DISCUSSION

Our studies of the significance of BCAA biosynthesis and transport by *B. anthracis* were prompted by data indicating that BCAA-related gene expression was altered by host-related signals and the virulence regulator AtxA. We were also intrigued by the unusually large number of *brnQ* genes associated with BCAA transport in *B. anthracis* as compared to most other firmicutes. We have determined that although *B. anthracis* can synthesize BCAAs, BCAA transport is required for optimal growth in culture and for virulence in a mouse model for anthrax. Moreover, we have obtained evidence for some functional redundancy among the multiple BrnQ transporters and identified major transporters of isoleucine and valine.

Other pathogenic *Bacillus* species, including *B. cereus* and *B. thuringiensis*, also possess an abundance of *brnQ* genes. *B. anthracis, B. cereus*, and *B. thuringiensis* have a large degree of chromosome gene synteny and share multiple physiological processes, but the three species have vastly different pathogenic lifestyles (52). While *B. anthracis* is capable of causing lethal systemic infections in mammals, certain strains of *B. cereus* that produce emetic and enterotoxins are causative agents of food poisoning and extraintestinal disease (53, 54), and some strains of *B. thuringiensis* are insect pathogens (55). Studies of BCAA synthesis and metabolism have not been reported for *B. cereus* and *B. thuringiensis*, but the presence of at least six putative BrnQ BCAA transporters in each species is intriguing given their differences in host interactions.

The apparent BCAA auxotrophy of *B. anthracis* despite the presence of BCAA biosynthesis genes is puzzling but not unprecedented among firmicutes. *L. monocytogenes* and *S. aureus* also carry genes that produce BCAA biosynthesis enzymes, but exhibit auxotrophic ambiguity (24, 27). The molecular mechanism(s) for this phenotype are not clear. In these species, BCAA biosynthesis is subject to multiple layers of control, including attenuation and CodY-mediated repression (27, 56). CodY, first identified in *B. subtilis* as a repressor of the dipeptide transport (*dpp*) operon (57), uses BCAAs and GTP as effectors to enhance DNA-binding activity at consensus sequences. CodY homologues in many Gram-positive bacteria have been reported to control a wide range of genes, including some virulence factors.

In *B. anthracis*, putative CodY-binding sites appear upstream of *ilvE1*, *ilvB2* and also in the promoter regions of *brnQ2*, *brnQ4*, *brnQ6*, and the BCAA-ABC transporter gene GBAA_1931 (58), but CodY-mediated regulation of these genes has not been reported. Interestingly, reports of CodY function in *B. anthracis* indicate an indirect role for CodY in toxin and capsule gene expression via AtxA. CodY does not affect transcription or translation of *atxA*. Rather, the regulator controls AtxA protein stability by an unknown mechanism (59). Our experiments examining *atxA* promoter activity and AtxA protein levels in cultures containing increasing levels of BCAAs showed no change in AtxA expression or protein level in response to BCAAs. Rather, our data revealed that BCAAs, most specifically valine, increase AtxA activity in a dose-dependent manner. Thus, BCAAs appear to be a signal for virulence gene expression in *B. anthracis*.

The mechanism for BCAA-enhanced AtxA activity is not known, but we surmise that it is CodY-independent. Van Schaik and coworkers (59) reported reduced AtxA protein levels in a *codY-*null mutant. We have been unable to construct a *codY-*null mutant in our parent strain. Yet, our experiments demonstrating BCAA activation of AtxA, taken together with our previous reports showing negative regulation of BCAA-related genes by XrrA (4) and positive regulation of XrrA by AtxA (15) suggest a feedback loop in which AtxA controls its own activity by regulating BCAA levels in cells. Although we did not find appreciable differences in the BCAA content of different mouse tissues, it is possible that niche-specific differences in BCAA availability could serve to modulate AtxA activity for optimal pathogenesis. Further investigations of relationships between BCAAs and AtxA are needed to dissect the molecular mechanism for this apparent fine-tuning of AtxA activity.

Our experiments examining growth and BCAA transport by mutants deleted for individual *brnQ* genes revealed that no single BrnQ transporter is essential for *B. anthracis* growth in our culture conditions. These findings are consistent with functional redundancy among the transporters. While none of the single *brnQ*-null mutants displayed growth defects in our culture conditions, some of the mutants were affected for BCAA transport, and complementation of mutants by inducing expression of the corresponding gene *in trans* resulted in elevated transport of some BCAAs. For example, deletion of *brnQ4* significantly decreased isoleucine and valine uptake, suggesting that BrnQ4 plays predominant role in transport of these BCAAs. A smaller reduction in isoleucine and valine uptake was observed for a *brnQ3*-null mutant. Interestingly, the uptake deficiency of the *brnQ3*-null mutant was fully restored by expressing *brnQ3 in trans*, but expression of *brnQ4 in trans* in the *brnQ4*-null mutant resulted in only partial restoration of isoleucine and valine transport.

We were unable to delete all *brnQ* genes in a single strain, indicating that at least one *brnQ* gene is essential for growth in culture. To test for specific function, we expressed individual *brnQ* genes *in trans* in a mutant deleted for *ilvD* and five of the six *brnQ* genes. We chose to create the *ΔilvDΔbrnQ2-6* mutant harboring an intact *brnQ1* gene for these studies based on BCAA uptake by single mutants and previously reported transcriptomic analyses. Expression of *brnQ1* is not affected during growth in blood and it is not controlled by AtxA (15, 34). The single *brnQ1*-null mutant showed a weak transport phenotype compared to other *brnQ*- null mutants (a small reduction in isoleucine and valine uptake) but the phenotype was not complemented by expressing *brnQ1 in trans*. A similar phenotype was associated with the single *brnQ5*-null mutant, but reduced isoleucine and valine uptake was recovered when *brnQ5* was expressed *in trans*. All other *brnQ* genes were of interest because they were highly regulated and/or single null mutants exhibited relatively strong transport phenotypes that could be complemented. The mutant, *ΔilvDΔbrnQ2-6*, was viable in our culture conditions but exhibited a significantly reduced growth rate, affirming the importance of BrnQ transporters for *B. anthracis* physiology. Further, the mutant showed a significant deficiency for isoleucine and valine uptake, suggesting that BrnQ1 does not play the major role in transport of these BCAAs. Data from our BCAA transport assays employing the *ΔilvDΔbrnQ2-6* mutant expressing individual BrnQ transporters revealed significant functional overlap for BrnQ3, BrnQ4 and BrnQ5 and strains expressing each of these transporters did not show significant differences in the kinetics of isoleucine and valine transport.

Assessment of leucine uptake by BrnQ3, BrnQ4 and BrnQ5 was not feasible because the *ΔilvDΔbrnQ2-6* mutant is capable of transporting leucine under our experimental conditions. Leucine transport may be associated with BrnQ1 or other uncharacterized non-BrnQ-type transporters. The *B. anthracis* genome sequence indicates a locus, GBAA_1931, _1933, _1934, _1935, and _1936, predicted to encode a BCAA-associated ABC transporter. Major leucine transporters have not been identified in other *Bacillus* species. However, a putative BCAA permease of *B. subtilis*, YvbW, has been reported to be regulated by a leucine-specific T-box mechanism and is a candidate for a leucine transporter (29, 60). We did not find a homologue of YvbW in the *B. anthracis* genome. A leucine-specific T box is present upstream of *brnQ6* (60), yet in our experiments leucine-uptake is unaffected in a *brnQ6*-null mutant.

Assignment of BCAA-specific function to BrnQ proteins is challenging not only because of functional redundancy, but because the regulation of the transporter genes is likely very complex. Experiments employing null-mutants and strains over-expressing *brnQ* genes in other bacteria have resulted in surprising phenotypes. For example, in *S. aureus*, deletion of *brnQ2*, encoding an isoleucine-specific transporter, affects the expression of *brnQ1* which encodes the major transporter for isoleucine, leucine, and valine (21). Expression of *brnQ1* is over 40-fold higher in a *brnQ2*-null mutant compared to the parent strain. Overexpression of *brnQ1*, results in elevated transport of isoleucine, leucine, and valine (21). Our *B. anthracis* data reveal the possibility of similar relationships in which one transporter affects the expression or function of another. The *brnQ3*-null mutant exhibited increased leucine uptake, a phenotype that was not complemented by expression of *brnQ3* from a xylose-inducible promoter *in trans*. It is possible that deletion of *brnQ3* elevated expression of a leucine transporter. In addition, growth of the *ΔilvDΔbrnQ2-6* mutant in the absence of BCAAs, although slow compared to parent, and a low rate of transport of isoleucine and valine by the mutant, indicate the presence of other additional transporters of these BCAAs. In this mutant, BrnQ1 or a non-*brnQ* type transporter may compensate for the absence of BrnQ3, BrnQ4, and BrnQ5.

Expression and function of BCAA transporters in the context of the host may differ from expression and function during culture. In our mouse model for systemic anthrax, *brnQ3* is essential for infection, whereas the *brnQ4*- and *brnQ5*-null mutants were fully virulent. BrnQ3- dependent virulence suggests that access to isoleucine and/or valine is critical for growth during infection, and that BrnQ3 serves as the major transporter of these BCAAs. It is possible, but less likely, that the absence of *brnQ3* alters expression of other transporters that are important for virulence. The importance of BCAA transport during infection has been reported for *S. aureus* and *S. pneumoniae* (16, 21, 22). In *S. aureus*, both BrnQ1 and BcaP are required for full virulence in a murine nasal-colonization and hematogenous spread infection model (16, 21). The *S. pneumoniae* branched-chain amino acid ABC transporter LivJHMGF is essential for virulence in a murine pneumonia model, but is not necessary for nasopharyngeal colonization (22).

The attenuated phenotype of the *brnQ3*-null mutant suggests limited availability of BCAAs in mouse tissues, resulting in growth restriction and/or reduced AtxA activity. Our LC-MS data revealed 0.08-0.23 nmol isoleucine mg^-1^ tissue, 0.15-0.41 nmol leucine mg^-1^ tissue, and 0.20-0.54 nmol valine mg^-1^ tissue. It is difficult to compare concentrations of BCAAs in solid tissues to those present in liquid, but it is worth noting that BCAA levels in human blood range from 0.02 mM to 0.27 mM and BCAA concentrations in porcine plasma are between 0.08 mM to 0.20 mM (18, 61, 62). Notably, in human blood and porcine plasma, as was true for our murine tissue samples, isoleucine is the least prevalent BCAA. Considering that the minimum concentration of BCAAs required for robust *B. anthracis* growth in culture is 0.25 mM, it is not unreasonable to consider these host niches as BCAA-limited.

Restricted availability of BCAAs in the animal is further supported by the inability of the *ilvD*-null mutant to establish infection in our animal model. Deletion of BCAA biosynthesis genes in other bacteria harboring apparent BCAA transporter genes, including *L. monocytogenes and S. pneumoniae*, also results in reduced virulence (20, 63). We note that while this manuscript was in review, Jenlinski et al. published work indicating that an *ilvD*-null mutant of *B. anthracis* strain 34F2 was not attenuated in a murine model using intranasal infection with spores (64). The bacterial strain and infection models differed from those employed in our study and the basis for this conflicting result will be examined in future studies.

Taken together, our data show that both BCAA transport and synthesis are critical for *B. anthracis* virulence in a murine model for late-stage anthrax. Further investigations will be directed at determining control mechanisms for BCAA-related genes and potential roles of BCAAs as host-related signals relevant for pathogenesis.

## MATERIALS AND METHODS

### Strains and growth conditions

Bacterial strains and plasmids used in this study are described in Table 1. *B. anthracis* strain ANR-1 (Ames non-reverting) was used as the parent strain. *Escherichia coli* TG1 was used for regular cloning and transformation. *E. coli* strains GM2163 and SCS110 (*dam^-^dcm^-^*) were used to isolate non-methylated plasmid DNA for electroporation of *B. anthracis* (65).

*E. coli* strains were cultured in Luria-Bertani (LB) broth (66) with shaking (150 rpm) or on LB agar plates at 37°C, with the exception of strains carrying the temperature-sensitive pHY304 vector which were cultured at 30°C. LB agar was also used for plating *B. anthracis.* When appropriate, antibiotics were used at the following concentrations: spectinomycin (50 µg ml^-1^ for *E. coli* and 100 µg ml^-1^ for *B. anthracis*), erythromycin (150 µg ml^-1^ for *E. coli* and 5 µg ml^-1^ for *B. anthracis*), and chloramphenicol (7.5 µg ml^-1^ for both *E. coli* and *B. anthracis*).

*B. anthracis* strains were pre-cultured overnight at 30°C in 25 ml of Brain Heart Infusion (BHI) medium (Becton, Dickson and Company, Franklin Lakes, NJ, USA) in a 250-ml flask with shaking (180 rpm). For each strain, 5 ml of overnight culture was centrifuged at 4000 rpm for 5 min, washed twice with phosphate-buffered saline (PBS) and then resuspended in PBS. The resuspended cells were transferred to a starting OD_600_ of 0.08 in 25 ml of Casamino Acids medium supplemented 0.8% sodium bicarbonate (CA-CO_3_) (44, 49) or in modified Ristroph medium (R) (10, 11). Unless indicated otherwise, the BCAA concentrations in R medium were as published (11): 1.75 mM isoleucine, 1.5 mM leucine, and 1.35 mM valine. Cultures were further incubated at 37°C with shaking in presence of 5% atmospheric CO_2_ (toxin-inducing conditions) (67) until the desired time-point or OD_600_ was reached. Growth curves for *B. anthracis* strains were performed in a 24-well sterile covered microplate using a Synergy HT Biotek plate-reader. The plate was incubated at 37°C in 5% atmospheric CO_2_ with continuous orbital shaking (365 cpm) and OD_600_ was determined at 1-h intervals.

### Recombinant DNA techniques

*B. anthracis* and *E. coli* were cultured overnight in 2 ml of LB broth. Genomic DNA was obtained from the *B. anthracis* cultures using the Ultraclean Microbial DNA isolation kit (Mo Bio Laboratories, Inc., Carlsbad, CA). Plasmid DNA isolation from *E. coli* was performed using the QIAprep Spin Miniprep Kit (QIAGEN Inc, Valencia, CA, USA). Oligonucleotides were purchased from Sigma-Aldrich (St. Louis, MO, USA) or Integrated DNA Technologies, Inc. (Coralville, IA, USA). For DNA cloning, Phusion HF DNA polymerase, high fidelity restriction enzymes, and T4 DNA ligase were purchased from either New England Biolabs (NEB) (Ipswich, MA, USA) or Thermo scientific (Waltham, MA, USA). The QIAquick Gel Extraction Kit and the QIAquick PCR Purification Kit (QIAGEN Inc, Valencia, CA, USA) were used for PCR-product clean-up. All procedures were performed according to the manufacturer’s protocol. Colonies of potential recombinant mutants were screened using colony PCR with 2.0 X Taq RED Master Mix (Genesee Scientific, USA). For colony PCR, a small quantity of cells was added to the PCR master mix and the suspension was heated at 95°C for 15 min prior to PCR. DNA sequences were confirmed by sequencing.

### Recombinant strain construction

Markerless gene deletions in *B. anthracis* were made by homologous recombination as described previously (5). Briefly, DNA fragments corresponding to sequences approximately 1 kb upstream and 1 kb downstream of target genes were obtained using PCR with the corresponding oligonucleotide pairs (see Table S1). Overlapping PCR was performed to obtain approximately 2-kb fragments representing the ligated flanking regions (68). Fragment preparations digested with appropriate restriction enzymes were ligated into plasmid pHY304, which contains a heat- sensitive origin of replication and an erythromycin-resistance cassette. Ligation mixtures were first transformed into *E. coli* TG1 (Table 1). Recombinant plasmids purified from TG1 were subsequently transformed into a *dam^-^dcm^-^ E. coli* strain to obtain plasmid DNA for electroporation of *B. anthracis*. *B. anthracis* strains were electroporated using a method described previously (5). Electroporants were passaged and plated on LB agar plates (master plates). Colonies from master plates were replica plated to LB agar and LB agar containing erythromycin to identify erythromycin-sensitive colonies. Sensitivity of erythromycin was indicative of the loss of the pHY304 construct with a potential double-crossover event resulting in recombination of the cloned DNA into the chromosome. Gene deletions were confirmed using PCR and sequencing. For creation of the *ΔilvDΔbrnQ2-6* mutant, the *ilvD, brnQ2, brnQ3, brnQ4, brnQ5 and brnQ6* genes were deleted sequentially in multiple experiments using distinct pHY304-derived constructs. Newly-constructed strains were stored as spores using a previously described protocol (69).

### Expression of transporter proteins

DNA corresponding to the sequences of the *B. anthracis brnQ1, brnQ2*, *brnQ3*, *brnQ4*, *brnQ5,* and *brnQ6* open reading frames was amplified from ANR-1 genomic DNA using PCR with the appropriate oligonucleotides as shown in Table S1. To facilitate detection of proteins encoded by these genes, amplification products were also generated carrying a sequence encoding a FLAG- tag fused to the 3’ end of the gene. The purified PCR products were digested with *Sal*I and *BamH*I restriction enzymes and finally ligated into the shuttle vector pAW285 such that expression was dependent on the xylose-inducible promoter (70). As described above, recombinant plasmids were first generated in *E. coli* TG1 (Table 1), and subsequently transformed into *dam^-^dcm^-^ E. coli* to obtain unmethylated plasmid DNA for electroporation into *B. anthracis*. For expression of the cloned genes, *B. anthracis* cultures were grown in CA-CO_3_ to OD_600_ ∼0.3, and induced with 1% xylose. After incubation for 3 h, cells were collected and cell lysates were prepared as described previously (5). Production of FLAG-tagged transporter proteins was assessed by western blotting using anti-FLAG antibody (Genscript, Piscataway, NJ, USA) as described previously (5).

### BCAA transport assay

Cells from cultures induced for transporter gene expression were collected using a Nalgene Rapid-Flow Sterile Vacuum Filter Unit (0.2 µM pore size) (Thermo fisher, Rockford, IL, USA). Cells were washed twice with PBS, resuspended in R medium lacking BCAAs to an OD_600_ of 5.0, and kept on ice until use. A 100-µl aliquot of prepared cells was stored at -20°C for measuring the protein content. The transport assay protocol was adapted from a previously published protocol (21, 32). Immediately before the assay, cells were incubated for 15 min at 37°C. The ^14^C-labeled amino acid of interest (Perkin Elmer, MA, USA) was added to the cells to a final concentration of 1 µM with stirring. At time intervals of 0, 30, 60, 90 and 120 seconds, 100-µl aliquots were removed, added to 4 ml of ice-cold 0.1 M LiCl_2_, and filtered rapidly through Whatman GF/F, 25 mm glass microfiber filters (GE healthcare, Chicago, USA) using a 1225 Sampling Manifold filtration unit (EMD Millipore, Darmstadt, Germany) attached to a vacuum pump. The filters were further washed with 4 ml of ice-cold 0.1 M LiCl_2_. The washed filters were dried for 30 min under a heat-lamp and placed in a 20-ml Borosilicate Glass Scintillation vial containing 5 ml of Ultima Gold F liquid scintillation cocktail (Perkin Elmer, MA, USA). ^14^C radioactivity retained on the filters was quantified using an LS6500 scintillation system (Beckman Coulter Inc., CA, USA). The protein contents of 100-µl cell samples were measured using the Bio-rad Protein Assay kit (Bio-rad, CA, USA). Uptake rates were calculated from retained activity and total added activity and expressed in pmol amino acid µg^-1^ of protein.

For kinetic measurements, cells were prepared as described above and then incubated for 20 sec with 0.5 µM, 1 µM, 2 µM, 4 µM or 8 µM of the radiolabeled substrate. Reactions were stopped by adding 4 ml of 0.1 mM ice-cold LiCl_2_ and cells was collected immediately using filtration. The initial velocity of uptake for each substrate was plotted using GraphPad Prism software version 9. Experiments were performed in triplicate and K_m_ and V_max_ values were determined using non-linear kinetics with GraphPad Prism software 9.

### Measurement of AtxA activity, AtxA protein levels, and *atxA* transcription

To quantify *in vivo* AtxA activity, we used the markerless *atxA*-null *B. anthracis* reporter strain UT376 that harbors the AtxA-regulated transcriptional fusion P*lef*-*lacZ* and contains an IPTG-inducible hexa-His-tagged *atxA* allele (pUTE991) (Table 1). Cultures were grown in R medium with various BCAA concentrations as indicated for 3 h and induced with 30 µM of IPTG. After incubation for 4 h, the OD_600_ was determined. One-ml samples were used for β-galactosidase assay, and 4-ml aliquots were preserved to quantify the AtxA protein using immunoblotting. β-galactosidase assays were performed as described previously (39) and expressed in Miller units. Western blotting was performed as described (5). His-tagged AtxA was detected using anti-His antibody (GenScript USA Inc., Piscataway, NJ, USA). RNA polymerase subunit β was used as a loading control and detected using anti-RNAP antibody (Thermo fisher, Rockford, IL, USA). Densitometry was performed using the ImageJ software (71) and AtxA protein levels were normalized with RNA polymerase β protein.

To assess *atxA* promoter activity, we used a *B. anthracis* reporter strain containing a pHT304-18z-derived *atxA* promoter-*lacZ* fusion (pUTE839) (38) (Table 1). Following induction with IPTG, one-ml samples were collected for β-galactosidase assays. Additional one-ml samples were used for total protein determination using the Bio-rad Protein Assay kit (Bio-rad, CA, USA). β-galactosidase assays were performed as described previously (38). Promoter activity was expressed as ONP min^-1^ mg^-1^.

### Virulence studies

Animal protocols were reviewed and approved by The University of Texas Health Science Center Institutional Animal Care and Use Committee and performed using accepted veterinary standards. Female 7-8 weeks-old A/J mice were purchased from The Jackson Laboratory (Bar Harbor, ME) and maintained in a pathogen-free environment. Mice were housed 5 per cage and were allowed a period of 72 hours to acclimate to their surroundings prior to use in experiments.

Infections and determinations of CFU in tissue were performed as described previously (4). Briefly, Mice were sedated with 0.1 mg ml^-1^ acepromazine administered intraperitoneally. Sedated mice were infected with 100 µl of a vegetative cell suspension (∼10^5^ CFU) via tail-vein injection. Mice were monitored four times daily for 11 days. Animals died naturally or were sacrificed upon the appearance of disease symptoms. The lungs, kidneys, livers, and spleens of deceased or sacrificed mice were aseptically excised, weighed, and placed individually in 1 ml of sterile DPBS containing zirconia/silica beads of diameter 2.3 mm (BioSpec Products, Inc, Bartlesville, OK, USA). Organs were homogenized by bead-beating for one min and then placed on ice. Tissue lysates were plated on LB agar and incubated overnight at 37°C for CFU determination. Kaplan-Meier survival curves and other statistical analyses were generated with GraphPad Prism version 9.

### RNA isolation

*B. anthracis* cultures were grown in R medium or R medium with 0.25 mM BCAAs for 5 h. Total RNA was extracted from 10 ml of culture using the NucleoSpin RNA kit (Macherey-Nagel Inc., PA, USA) as described by the manufacturer. RNA samples were quantified using a NanoDrop ND-1000 Spectrophotometer. Isolated RNA was treated with RNase-Free DNase (QIAGEN Inc, Valencia, CA, USA) for 2 h at room temperature to remove any genomic DNA contamination. RNA was then purified and concentrated using RNA Clean And Concentrator Kit (Zymo Research, Irvine, CA, USA). Removal of DNA contamination was verified by PCR using purified RNA as template.

### Reverse Transcription (RT) PCR

Non-quantitative RT-PCR was used to test for co-transcription of the *ilv* genes. Purified RNA was used for the first strand cDNA synthesis employing the Superscript III Reverse transcriptase kit (Invitrogen, Carlsbad, CA, USA) and random hexamers. Synthesized cDNA was purified using a QIAquick PCR Purification Kit (QIAGEN Inc, Valencia, CA, USA) and the final cDNA concentration was determined using a NanoDrop. Purified cDNA (45ng) was used as template with primers shown in Table S1 to amplify transcripts extending across adjacent genes. The PCR products were examined in a 1.2% agarose gel after ethidium bromide staining.

### Branched chain amino acid measurement in mouse tissues

Lungs, livers, spleens, and kidneys were harvested from each of three uninfected 7- to 8-week-old female A/J mice, placed in 15-ml centrifuge tubes, and snap-chilled in liquid nitrogen. BCAA analysis was performed by the M.D. Anderson Metabolomics Core Facility (Houston, TX). A small portion of each tissue sample was weighed and transferred to a 2 ml Precellys tube containing ceramic beads (Bertin, France). Tissue samples (20 to 30 mg) were crushed in liquid nitrogen and homogenized using Precellys Tissue Homogenizer (Bertin, France). Amino acids were extracted using 0.5 ml of ice-cold 90/10 (v/v) methanol/water containing 0.1% formic acid. Extracts were centrifuged at 17,000 g for 5 min at 4°C, and supernatants were transferred to clean tubes, followed by evaporation to dryness under nitrogen. Samples were reconstituted in 0.1% formic acid in 90/10 (v/v) acetonitrile/water, then injected for analysis by liquid chromatography (LC)-MS. LC mobile phase A (MPA; weak) was acetonitrile containing 1% formic acid, and mobile phase B (MPB; strong) was water containing 50 mM ammonium formate. A Thermo Vanquish LC system included an Imtakt Intrada Amino Acid column (3 µm particle size, 150 x 2.1 mm) with column compartment kept at 30°C. The autosampler tray was chilled to 4°C. The mobile phase flow rate was 300 µL min^-1^, and the gradient elution program was: 0-5 min, 15% MPB; 5-20 min, 15-30% MPB; 20-30 min, 30-95% MPB; 30-40 min, 95% MPB; 40-41 min, 95-15% MPB; 41-50 min, 15% MPB. The total run time was 50 min. Data were acquired using a Thermo Orbitrap Fusion Tribrid Mass Spectrometer under ESI positive ionization mode at a resolution of 240,000. Raw data files were imported to Thermo Trace Finder software for final analysis. The relative abundance of each metabolite was normalized by tissue weight.

## ACKNOWLEDGEMENTS

This work was supported by the National Institute of Allergy and Infectious Diseases (NIAID) grants R01 AI33537 and R21 AI151313 to T.M.K. During a portion of this work N.B. was supported by the NIAID training grant T32 AI55449. S.D. and T.M.K. conceived of the project, designed the experiments, analyzed the data, and wrote the manuscript. I.D.C. made intellectual contributions, assisted with animal experiments, and edited the manuscript. N.B. made intellectual contributions. All authors read and approved the manuscript.

We are indebted to Dr. Heidi Vitrac, Ph.D., for helping us establish the BCAA transport assays. We thank Malik Raynor, Ph.D., and Jung-Hyeob Roh, Ph.D. for their helpful discussions of the work. The metabolomics facility at M.D. Anderson Cancer Center, Houston, Texas performed BCAA measurements in mouse tissues.

The content of this publication is solely the responsibility of the authors and does not necessarily represent the official views of the National Institute of Allergy and Infectious Diseases. We declare no competing interests.

---

**Table S1:** Oligonucleotides used in this study.

**Figure S1: Graphical representation of BCAAs in mouse tissues.** (A) Isoleucine, (B) Leucine, (C) Valine. One-way ANOVA followed by Tukey’s multiple comparison test was used for data analysis. Each bar represents the mean of three biological replicates. Error bars represent +/- standard deviation. Asterisks indicate P-values. (* =P<0.05, **=<0.01, *** = <0.001, **** =P<0.0001).

